# Genome-wide H3K9 Acetylation Level Increases with Age-Dependent Senescence of Flag Leaf in Rice (*Oryza sativa*)

**DOI:** 10.1101/2021.11.14.468555

**Authors:** Yu Zhang, Yanyun Li, Yuanyuan Zhang, Zeyu Zhang, Deyu Zhang, Xiaonan Wang, Binfan Lai, Dandan Huang, Lianfeng Gu, Yakun Xie, Ying Miao

## Abstract

Flag leaf senescence is an important biological process that drives the remobilization of nutrients to the growing organs of rice. Leaf senescence is controlled by genetic information via gene expression and epigenetic modification, but the precise mechanism is as of yet unclear. Here, we analyzed genome-wide acetylated lysine residue 9 of histone H3 (H3K9ac) enrichment by chromatin immunoprecipitation-sequencing (ChIP-seq) and examined its association with transcriptomes by RNA-seq during flag leaf aging in rice (*Oryza sativa*). We found that genome-wide H3K9 acetylation levels increased with age-dependent senescence in rice flag leaf, and there was a positive correlation between the density and breadth of H3K9ac and gene expression and transcript elongation. A set of 1,249 up-regulated, differentially expressed genes (DEGs) and 996 down-regulated DEGs showing a strong relationship between temporal changes in gene expression and gain/loss of H3K9ac was observed during rice flag leaf aging. We produced a landscape of H3K9 acetylation-modified gene expression targets that includes known senescence-associated genes, metabolism-related genes, as well as miRNA biosynthesis-related genes. Our findings reveal a complex regulatory network of metabolism- and senescence-related pathways mediated by H3K9ac and also elucidate patterns of H3K9ac-mediated regulation of gene expression during flag leaf aging in rice.

**Significance statement:** Genome-wide H3K9 acetylation levels increased with age-dependent senescence in rice flag leaf, and positively correlation the density and breadth of H3K9ac with transcript elongation and expression. Identified numerous H3K9 acetylation-modified gene expression targets reveal a complex regulatory network and metabolism-mediated senescence network that are associated with H3K9ac during leaf aging in rice.

## INTRODUCTION

Leaf senescence is the highly ordered final phase in leaf development. This process is characterized by the degradation of chlorophylls, nucleic acids, lipids, proteins, and other macromolecules, and subsequently by programmed cell death (Lim et al., 2007). As the nutrients are recycled to the developing parts of plants such as seeds or growing organs, leaf senescence therefore can be considered an important developmental process for plant fitness (Woo et al., 2019). However, in the agricultural setting, leaf senescence may limit crop yield by shortening the growth phase (Lim et al., 2007; Schippers et al., 2015). Understanding the regulatory mechanism of leaf senescence in rice is thus critical for the production of high yielding crops.

The transition into leaf senescence is preceded and accompanied by alteration of a series of gene expression. Global gene expression analyses revealed that dramatic changes in the expression of thousands of senescence-associated genes (SAGs) occur during flag leaf senescence in rice (Sato et al., 2011; Liu et al., 2016; Lee et al., 2017). A leaf senescence database currently compiles 188 SAGs experimentally identified in rice (Li et al., 2020). Among them, various TFs, including NAC (NAM, ATAF, CUC), WRKY, zinc finger, HD-Zip III (Class III Homeodomain-Leucine Zipper), and bHLH (Basic Helix-Loop-Helix) family proteins, play pivotal roles in the transcriptional regulation of SAGs during leaf senescence (Leng et al., 2017). However, the expression of these TFs is also regulated during leaf senescence, suggesting that additional mechanisms determine the onset of leaf senescence (Yolcu et al., 2018; Woo et al., 2019).

In addition to cis- and transacting factors that induce SAGs at the transcriptional level, recent findings have shed light on the epigenetic regulation of plant senescence through non-coding RNAs and histone modification (Yolcu et al., 2018; Ostrowska-Mazurek et al., 2020). Acetylated Lys residue 9 of histone H3 (H3K9ac) is one of the most extensively studied epigenetic markers in plants, and is known as a marker of actively transcribed chromatin (He et al., 2010; Zhou et al., 2010; Du et al., 2013). The correlation between H3K9ac with gene expression during de-etiolation and after UV-B treatment has been reported (Charron et al., 2009; Schenke et al., 2014). The timing of H3K9ac enrichment or loss closely parallels gene expression changes for the circadian clock event and response to drought stress (Kim et al., 2012; Malapeira et al., 2012), or it occurs after genes have been activated in the case of flowering (Adrian et al., 2010), and aging in Arabidopsis (Brusslan et al. 2015; Huang et al. 2018). However, the exact role of H3K9ac in gene activation is not yet defined. Especially in rice, it is unclear how the transcriptional reprogramming of senescence-related genes is regulated by H3K9ac modification in a developmental-dependent manner.

In this study, we analyzed genome-wide H3K9ac enrichment using ChIP-seq and examined its association with gene expression by RNA-seq at different stages of flag leaf aging in rice. Histone acetylatome analysis revealed that the density and breadth of H3K9ac positively correlated with gene expression. Genetics analysis revealed a landscape of H3K9ac-associated DEGs during rice flag leaf aging. These results uncover a complex regulatory network and metabolism-mediated senescence network associated with H3K9ac during leaf aging in rice.

## Materials and Methods

### Plant materials and growth condition

Rice (*Oryza sativa*) ssp. japonica (*Nipponbare*) plants were grown in a greenhouse under 16 h light/8 h dark cycle at 32 and 28°C, respectively. We used the flag leaves from the first tiller at 0, 1, 2, 3, 4 and 5 weeks after full heading for all experiments in this study. Leaves at the independent stages were harvested, frozen in liquid nitrogen, and stored at -80°C for RNA isolation or processed directly after harvesting for ChIP assay.

### Photosynthetic measurements

To detect the maximum photochemical efficiency of photosystem II (Fv/Fm), the chlorophyll fluorescence parameters from the middle zones of flag leaf were measured with IMAGING-PAM MAXI (Heinz Walz GmbH, Effeltrich, Germany) after dark-adaptation for 15 min as described by Shao (2007). For chlorophyll measurements, the middle zones of flag leaves were detected using a plant nitrogen balance index measuring device (DUALEX SCIENTIFIC+). Data from five plants were collected for each stage.

### ChIP assays

ChIP assays were performed according to a published protocol (Gendrel et al., 2005) with minor modification. Briefly, 1.5 g of pooled rice flag leaf tissue was harvested and cross-linked with 1% (v/v) formaldehyde under a vacuum for 15 min. Fixation was stopped by adding glycine at a final concentration of 0.125 M. The cross-linked tissue was ground into powder in liquid nitrogen, followed by nuclear isolation with the Plant Nuclei Isolation/Extraction Kit (Sigma-Aldrich) and sonication with a Bioruptor (Diagenode) to 200 bp of average fragment size. Protein A-agarose beads (Millipore) and an antibody against H3K9ac (Millipore, #07-352) were used to precipitate the DNA, which was digested by proteinase K (Ferments), recovered using a QIAquick PCR purification kit (Qiagen). The ChIP DNA and input DNA were recovered and dissolved in water for ChIP-seq and ChIP-qPCR analysis.

### Construction of ChIP-seq libraries and ChIP-qPCR analysis

For ChIP-Seq, flag leaf samples at weekly intervals from 0 to 5 weeks after full heading were harvested, and each stage of flag leaf was pooled from ten rice plants and three biological replicates were prepared for each input and IP. Chromatin from flag leaves were immunoprecipitated with an antibody against H3K9ac (Millipore, #07-352). ChIP-DNA and input DNA libraries were constructed and sequenced by Beijing Novogene Bio-Tech. ChIP DNA, pooled from three independent ChIP experiments, were end-repaired, followed by A-base addition and ligation with adapters using TruSeqPECluster Kit (Illumina). After PCR enrichment, the DNA library was quantified with Qubit (Thermo Fisher), followed by cBotcluster generation (Illumina). The libraries were sequenced on an Illumina Hiseq platform and 50 bp single reads were generated.

For ChIP-qPCR, ChIP DNA was analyzed by qRT-PCR using the primer pairs (Table S3). Fragments (2,000 bp) of sequences upstream of ATG were chosen as the promoter regions for primers design. All primers amplified 80-to 120-bp products and were annealed at 58°C. ChIP DNA enrichment was determined as % of input (i.e., the relative amount of immunoprecipitated DNA compared to 100% input DNA after qPCR analysis). Each PCR was performed with three technical replicates and three biological replicates.

### ChIP-seq data analysis

For ChIP-seq analysis, sequencing raw data were cleaned to filter low quality data, and the clean reads were aligned to the rice reference genome of the version 7 (MSU7.0) by BWA (version 0.7.12) with default parameters (Li, H. and R. Durbin, 2009). Peak calling was performed using MACS2 (version 2.1.0). The “–mfold” parameter was set as “5-50”and a q value threshold of enrichment of 0.05 was used for all data sets (Zhang et al., 2008). For viewing the data, the.bw files were visualized with the IGV genome browser (http://software.broadinstitute.org/software/igv/download). To obtain the distribution of the reads through the genes, each gene (including the 2-kb regions upstream and downstream of each gene and gene body regions) was divided into 100 intervals (1% each interval), and the read density was then calculated by: each interval read numbers/total read numbers per interval length. Different peak analyses were based on the fold enrichment of peaks of different experiments. A peak was determined as different peak when the odds ratio between two groups was more than 2. Using the same method, genes associated with different peaks were identified and GO and KEGG enrichment analysis performed. The visualization of the average read coverage over gene body and additional 2 kb upstream and downstream of the TSS and TES was performed by deepTools2 (Ramírez et al., 2016). The distribution of peak summits on different function regions, such as promoter, 5’ UTR, 3’UTR, exons, introns and intergenic regions was performed with the R package ChIPseeker (Yu et al., 2015).

### RNA-Seq and data analysis

For continuous gene expression profiling, the flag leaf was collected at weekly intervals from 0 to 5 weeks after full heading like above. Three independent biological replicates were performed for all leaf stages, each of them being a pool of flag leaves from ten individual plants. Total RNA was extracted with Trizol reagent (Invitrogen), followed by DNase I digestion (Promega). The quality of the total RNA was assessed using the Nanodrop spectrophotometer and Agilent 2100 Bioanalyzer. Eighteen RNA-seq libraries were constructed by the dUTP method according to the manufacturer’s instructions as described by Wang et al. (2017). Paired-end 150–bp reads were sequenced with the Illumina HiSeq X10 platform (Biomarker Biotechnologies).

The RNA-seq data were aligned against the TIGR7 genome using TopHat 2.0.11 (Trapnell et al., 2009), with default parameters. The gene expression levels were measured and normalized as FPKM (fragments per kilobase of transcript per million mapped reads). We used edgeR (Nikolayeva and Robinson, 2014) to call differentially expressed genes (DEGs) with |log2_ratio|≥1 and FDR≤0.01. GO enrichment was analyzed in AgriGO (Tian et al., 2017). The Yekutieli method was used for multiple test correction, and terms with a false discovery rate greater than 0.05 were discarded. GO terms were summarized using REVIGO (Supek et al., 2011). The results were then imported into Cytoscape for visualization. KEGG pathway enrichment analysis was done using KOBAS software (KOBAS, Surrey, UK). The statistical test method is hypergeometric test / Fisher’s exact test and the FDR correction method is Benjamini and Hochberg.

### Gene co-expression network analysis

The differential expressed genes were used to construct co-expression networks by the R package WGCNA (Langfelder and Horvath, 2008). The soft power parameter of 8 was used to derive the correlation matrix and optimize topology network. Spearman correlation coefficient was used to build adjacency matrices to calculate topographical overlap matrices (TOMs), which were constructed to calculate edge weights with default parameters. Finally, hierarchical clustering of TOM (1-TOM) distance was used to identify sub networks. Similar modules were merged using the merge Close Modules function with cut Height=0.2.

### RT-qPCR analysis

Total RNA was reverse transcribed using the TransScript® One-Step gDNA Removal and cDNA Synthesis SuperMix kits (TransGen Biotech). Primers for qRT-PCR are listed in Table S3. The reverse transcription products were used as templates for quantitative real-time PCR performed on a CFX96 real-time PCR system (Bio-Rad) using SYBR Green Master Mix (DBI Bioscience) according to the manufacturer’s protocol. For all reactions, *OsACTIN* (*LOC_Os11g06390*) was used as an internal control. Relative expression levels were measured using the 2^-ΔΔCt^ analysis method. All reactions were performed in triplicate with three independent experiments.

### Statistical analysis

Data are given as mean ± SD and were analyzed by Student’s t test or one-way ANOVA, and differences with P-values <0.05 or <0.01 or <0.001 were considered statistically significant.

## Data availability

ChIP-seq data and RNA-seq raw data are available from the Sequence Read Archive (https://www.ncbi.nlm.nih.gov/sra) under accession number SRP331485.

## RESULTS

### The H3K9ac profile changes during rice flag leaf aging

To characterize the leaf aging behavior of rice plants, the phenotypes of flag leaves at 0 to 5 weeks after heading (WAH), 77 days after germination, were observed. Leaf chlorophyll content, Fv/Fm-based photochemical efficiency of PSII and the expression patterns of several senescence-associated genes were measured at the indicated times to monitor the progression of leaf senescence. The flag leaves remained green from 0 to 2 WAH and started to show yellowing at the tip after 3 WAH (Figure S1A). Similarly, the Fv/Fm ratios of flag leaves were maintained at a similar level from 0 to 2 WAH, with a significant decrease after 3 WAH (Figure S1B). The chlorophyll content was first increased and started to drop considerably earlier at 1 WAH (Figure S1C). In addition, the expression of the senescence-associated genes *OsSGR* and *OsONAC011* were up-regulated significantly from 1 WAH, and both showed a tendency toward increased expression during aging (Figure S1D). These results show that the time period sampled encompassed the onset and progression of flag leaf senescence in rice.

To gain insight into the role of H3K9ac modification in rice leaf senescence, we conducted twelve ChIP-seq libraries in rice flag leaf tissues at the six time points mentioned above using an antibody against histone H3K9ac. Approximately 71% to 84% of the ChIP-seq reads were mapped to unique positions in the rice genome sequence (TIGR7) during leaf aging (Table S1). The distribution of H3K9ac markers in the genome showed that H3K9ac peaks were predominantly enriched in the intragenic and promoter regions, which occupied a major part of the H3K9ac peaks (Figure 1A; Table S2). Further inspection of the average read coverage in different genomic regions of genes revealed that H3K9ac modification formed strong peaks within the region from transcription start sites (TSS) to 240 bp downstream during leaf aging (Figure 1B). In general, this distribution pattern around TSS is very similar to the previous report in *Arabidopsis* (Brusslan et al., 2015). The conserved H3K9ac distribution pattern at the TSSs of rice and *Arabidopsis* suggests that H3K9ac plays a significant role in gene transcription regulation. It is worth noting that H3K9ac levels increased with the transition of the leaves from 0 WAH to 5 WAH (Figure 1B).

**Figure 1.**
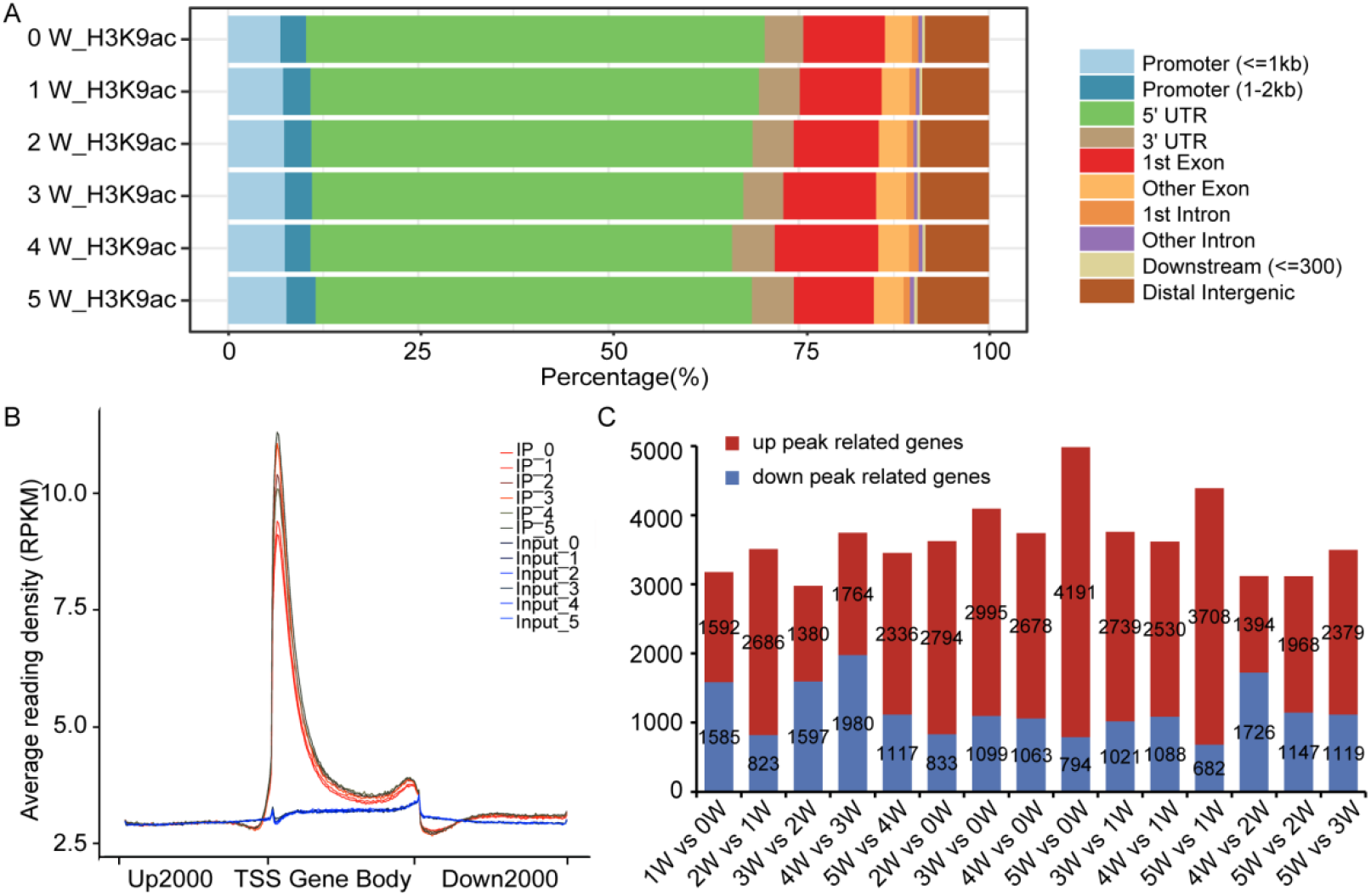
Genome-wide mapping of H3K9ac sites across rice flag leaf development. (A) Stacked bar plot showing the distribution of H3K9ac peaks in different genomic regions from 0 to 5 WAH. Promoters are defined as genomic regions encompassing 2 kb upstream of the TSS. Intragenic regions contain 5’ UTR, exons, introns, and 3’ UTR. Downstream (<=300) means 300 bp downstream of the transcription end site. Distal intergenic regions are genomic regions except downstream (<=300) and promoter regions. (B) Characterization of the H3K9ac distribution pattern for genes at six stages of leaf aging. The x axis represents different regions of a gene, and the y axis represents the average read density normalized by sequencing depth and window size of 20 bp (RPKM). The peaks of read counts are most commonly located in the 240-bp region downstream of the TSS. (C) Numbers of H3K9ac peak-related genes. Red indicates gain peak-related genes, and blue indicates loss peak-related genes.

To investigate whether levels of the H3K9ac marker changed significantly during flag leaf aging, comparisons were made between six time points. A gene was defined as a differentially H3K9ac-modified gene (DHG) if the peak enrichment (IP/Input) fold change between different time points was >2 and FDR <0.05. We found 10,996 differentially H3K9ac-modified genes including 9,272 genes with increased H3K9ac modification and 5,903 genes with decreased H3K9ac modification (Figure 1C). A larger number of genes with increased H3K9ac modification were observed overall, suggesting a global increase of H3K9ac modification with rice flag leaf aging.

GO enrichment analysis of the diff-peak-related genes showed that the genes associated with a loss of H3K9ac modification were enriched in various biological processes, including photosynthesis (light reaction, photosystem II assembly), sulfur compound biosynthetic processes, small molecule biosynthetic processes, cysteine biosynthetic processes and metabolic processes during rice flag leaf aging. Genes associated with an increase in H3K9ac were enriched for L-ascorbate peroxidase activity in 1 vs 0 WAH comparison. Genes related to starch metabolic processes and transferase activity were enriched in 2 vs 1 WAH and 4 vs 3 WAH comparisons. In 5 vs 0 WAH and 5 vs 1 WAH comparisons, the gain diff-peak-related genes were enriched for transmembrane signaling receptor activity and catalytic activity (Figure 2).

**Figure 2.**
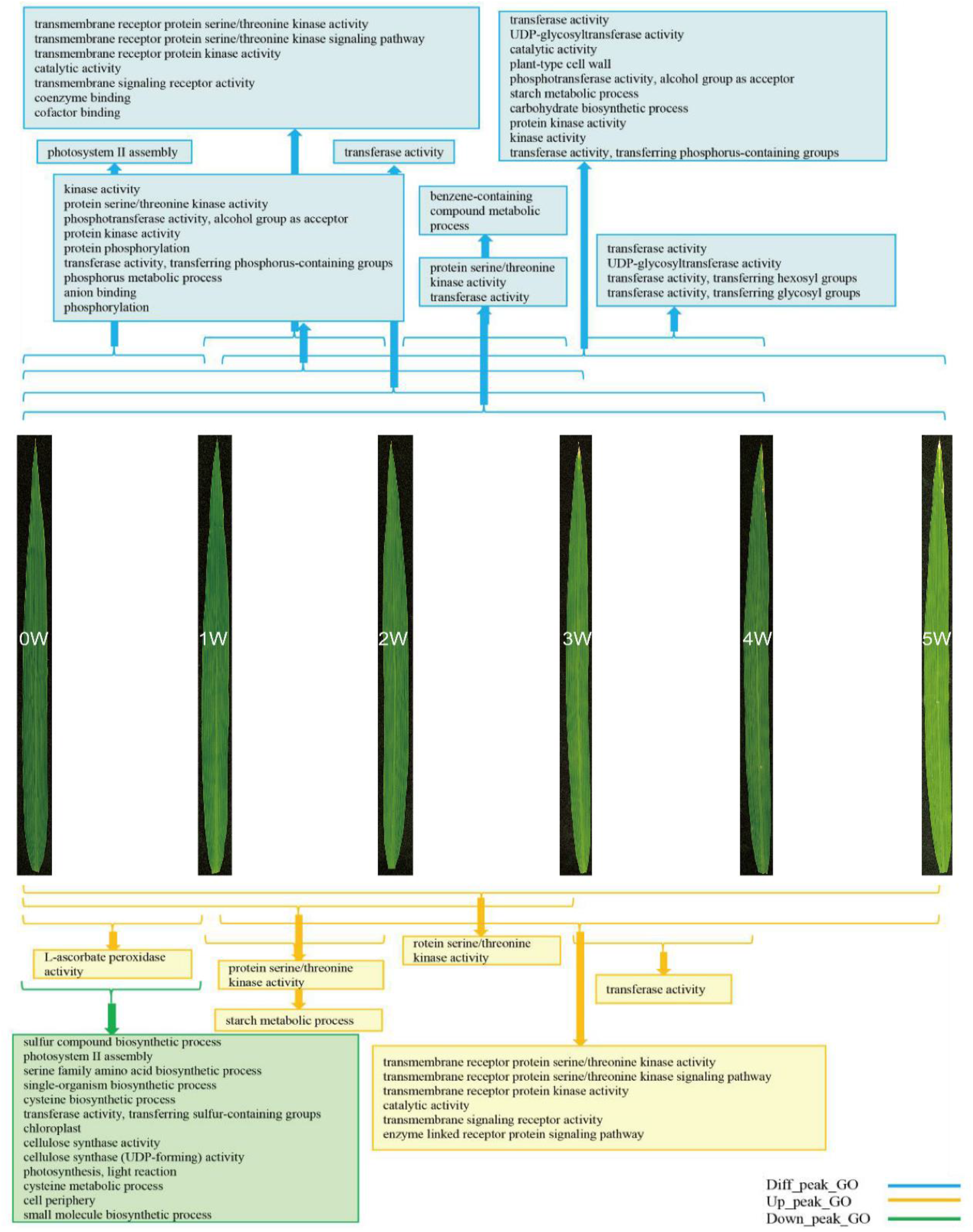
Biological processes promoted or repressed at different time points during senescence. Enriched GO terms were identified in groups of genes that show significant up-regulation or down-regulation H3K9ac modification during leaf development and senescence. White numbers indicate WAH. Yellow pale is enriched in gain-H3K9ac peak-related genes; green pale is enriched in lose-H3Kac peak-related genes; blue pale is enriched in all differential H3K9ac-modified genes.

Interestingly, Kyoto Encyclopedia of Genes and Genomes (KEGG) pathway analysis showed that the loss diff-peak-related genes were mainly involved in glutathione metabolism, steroid biosynthesis, ascorbate and aldonate metabolism, porphyrin and chlorophyll metabolism pathways in 2 vs 1 WAH and 2 vs 0 WAH comparisons. The most abundant gain diff-peak-related genes were related to plant-pathogen interaction, lipid and carbohydrate metabolism, zeatin biosynthesis and brassinosteroid biosynthesis pathways in 3 vs 0 WAH, 3 vs 2 WAH and 5 vs 2 WAH comparisons (Table 1). The KEGG enriched pathway results are consistent with the metabolism processes of rice flag leaf senescence (Lim et al., 2007; Woo et al., 2019), suggesting that H3K9ac modification profile is closely associated with flag leaf senescence in rice.

**Table 1.** The considerably changed KEGG pathways for differential H3K9ac-modified genes during rice developmental flag leaf aging. Over-represented enriched KEGG pathways (p-value<0.05) are reported.

|  | Comparisons | Pathways | Gene Count | p-Value |
| --- | --- | --- | --- | --- |
| UP_KEGG<br>Pathway: | 1W VS 0W | Ascorbate and aldarate metabolism | 8 | 1.23E-03 |
|  |  | Phenylpropanoid biosynthesis | 19 | 3.99E-03 |
|  | 3W VS 0W | Plant-pathogen interaction | 25 | 6.81E-04 |
|  |  | Zeatin biosynthesis | 7 | 2.67E-03 |
|  | 3W VS 2W | alpha-Linolenic acid metabolism | 7 | 1.37E-03 |
|  |  | Brassinosteroid biosynthesis | 4 | 1.46E-03 |
|  |  | Phenylalanine metabolism | 13 | 3.28E-03 |
|  |  | Phenylpropanoid biosynthesis | 15 | 5.70E-03 |
|  | 5W VS 2W | Glycolysis / Gluconeogenesis | 17 | 1.92E-03 |
| Down_KEGG<br>Pathway: | 2W VS 0W | Glutathione metabolism | 8 | 1.65E-03 |
|  |  | Steroid biosynthesis | 5 | 4.59E-03 |
|  | 2W VS 1W | Ascorbate and aldarate metabolism | 6 | 8.54E-04 |
|  |  | Porphyrin and chlorophyll metabolism | 5 | 4.31E-03 |
|  | 4W VS 0W | Galactose metabolism | 7 | 3.45E-03 |
|  |  | Starch and sucrose metabolism | 13 | 4.32E-03 |

### Time course transcriptomes during flag leaf aging

H3K9ac is one of the most extensively studied epigenetic markers in plants, and is also known as a marker of actively transcribed chromatin (He et al., 2010; Du et al., 2013). The enrichment of H3K9ac at the TSS region promoted gene expression at the development stages of the *Arabidopsis* plant (Brusslan et al. 2015; Huang et al. 2018). To study the potential role of H3K9ac in gene expression during rice flag leaf aging, we chronologically performed RNA-seq analysis of the above-mentioned plant materials. To investigate the gene activity dynamics during rice leaf aging, we compared expression in all 15 pairs of the six different time points. This analysis revealed that in total, 5,694 genes were differentially expressed (DEGs, |Log2 fold change| > 1 and FDR < 0.01), including 3,018 up-regulated and 3,155 down-regulated DEGs (Figure 3A). Adjacent time point comparisons analysis showed that the largest number of genes was up- or down-regulated during the first-time interval, suggesting that major changes in gene expression occur before the visual manifestations of leaf aging: the loss of chlorophyll.

**Figure 3.**
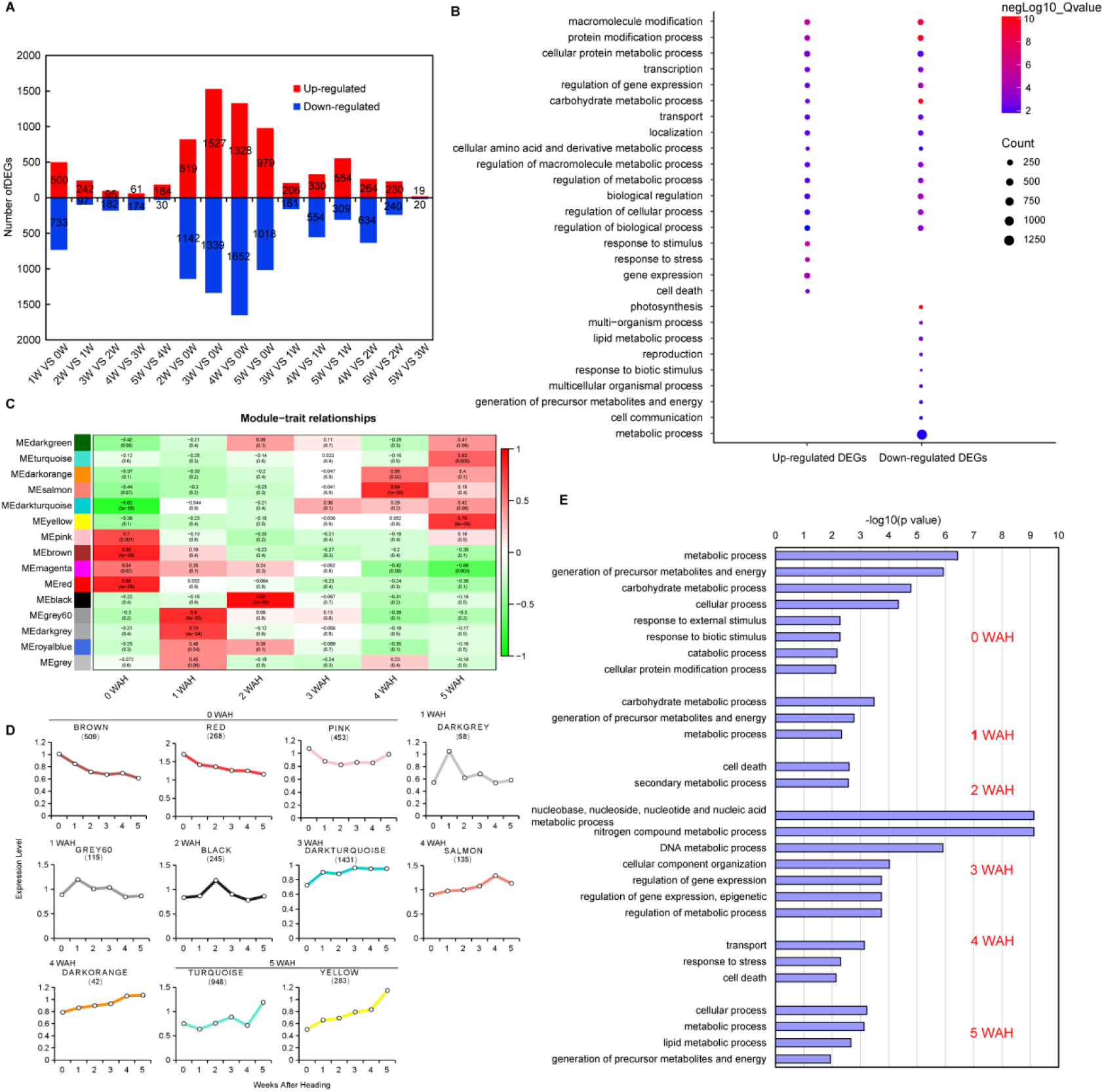
Gene expression differences during rice flag leaf aging. **(A)** The numbers of genes showing differential expression (≥2-fold, FDR≤0.01) in all 15 pairs of the six different time points. **(B)** Gene ontology biological processes (GOBPs) significantly (Q < 0.05) represented by up- and down-regulated DEGs. The color bar denotes the gradient of -log 10 (Q) values. **(C)** Module-samples associations. Weighted gene co-expression network analysis (WGCNA) of differentially-expressed genes (DEGs) identified from rice flag leaves over six aging stages. The left panel shows the fifteen modules and each row corresponds to a module. The color of each cell at the row-column intersection indicates the correlation coefficient between the module and the sample type. A high degree of positive correlation between a specific module and the sample type is indicated with dark red. Negative correlation is indicated with green. The numbers in parentheses are the corresponding P-values. **(D)** Gene expression patterns of stage-specific modules during rice flag leaf aging. Each graph shows the expression pattern of the module. The y-axis indicates the mean log10 (FPKM+1) of all genes within a given module. The x-axis indicates the stages. The number of genes residing in each module is given in parentheses. **(E)** Biological processes enriched in sample-specific modules. Column length indicates the overrepresentation significance of a group of GO terms.

GO enrichment analysis was performed to explore the functional significance of DEGs. It revealed 14 biological processes (GO terms) to be commonly enriched in up- and down-regulated DEGs. Genes related to transcription activity, defense and cell death were most significantly enriched in the up-regulated genes. Genes related to photosynthesis and growth-related processes, such as multi-organism process, reproduction, lipid metabolic processes, and cell communication, were most significantly enriched in the down-regulated genes (Figure 3B).

To further provide insights into the functional transitions along leaf aging, we clustered 5,694 DEGs into 15 co-expression modules using the weighted gene co-expression network analysis (WGCNA), and then performed functional analysis for each stage-specific module. We identified 15 modules that were color-coded and ranged in size from 42 genes (dark orange module) to 1,431 genes (dark turquoise module; Figure 3C). The module with a correlation coefficient above 0.63 and p < 0.05 was defined as the sample-specific module. Thus, ten tissue-specific modules were obtained, as shown in Figure 3C. Brown, red and pink modules were positively correlated with 0 WAH; Dark grey and Grey were positively correlated with 1 WAH; Black was positively correlated with 2 WAH; the specific modules of 4 WAH were Salmon and Dark orange; the specific modules of 5 WAH were Turquoise and Yellow (Figure 3C). Because the dark turquoise module with a correlation coefficient of 0.35 and p < 0.1 was not a sample-specific module, to identify genes positively correlated with this time point, a set of 395 DEGs with expression levels that peaked at 3 WAH were screened from the dark turquoise module manually (Supplementary Dataset S2). The expression profiles of the respective modules are depicted in Figure 3D. Genes in a module have preferential expression in a particular stage relative to all other samples, further confirming the positive correlation between the modules with the leaf stage.

Gene ontology enrichment analysis of stage-specific modules highlighted key biological processes during flag leaf aging. Modules associated with early stages of development (0 and 1 WAH) showed enrichment of GO terms related to generation of precursor metabolites and energy, and carbohydrate metabolic processes; GO term related to cell death was represented at 2 WAH, including two expression protein genes (*LOC_Os10g04570, LOC_Os04g02030*) and one resistance protein gene *RGA2* (*LOC_Os05g41310*), and at 4 WAH, including *LOC_Os09g14490* and *LOC_Os10g33440*; The 3 WAH stage-specific modules represented GO terms were related to nitrogen compound metabolic processes, DNA metabolic processes, epigenetic regulation of gene expression and cellular component organization. The epigenetic regulation of gene expression GO term includes *SDG714* encoding a histone H3K9-specific methyltransferase, *WAVY LEAF1* encoding an RNA methyltransferase, *OsAGO1d, CROWN ROOT DEFECT 1, LOC_Os04g54840* encoding a *DNA-directed RNA polymerase subunit, LOC_Os03g22570* encoding MIF4G domain-containing protein, as well as two HhH-GPD superfamily base excision DNA repair genes. The modules associated with late development (4 and 5 WAH) exhibited overrepresentation of GO terms, such as transport, response to stress, cell death and lipid metabolic processes, etc. (Figure 3E). Further inspection showed that *Oryza sativa* monosaccharide transporters 3 (*OsMST3*; *LOC_Os07g01560*), peptide transporter 7 (*OsPTR7*; *LOC_Os01g04950*) and a gene encoding a protease inhibitor *LOC_Os11g02389* were up-regulated in the late stage of leaf aging. These results suggested that each of the leaf development stages was associated with one or more co-expression modules that reflected the existing gene regulatory processes corresponding to each stage during flag leaf aging.

### The density and breadth of H3K9ac correlates with gene expression during flag leaf aging

To obtain a comprehensive understanding of global interaction patterns between transcription and epigenetic modification, we generated an integrated epigenomic map by sorting all genes with H3K9ac modification based on their expression levels (Figure 4A). A coordinated regulation of both H3K9ac enrichment and gene activation was observed, in which genes with low levels of H3K9ac enrichment showed low levels of expression, while genes with high levels of H3K9ac enrichment exhibited high levels of expression; these were mainly positioned after the TSSs (Figure 4A). After dividing all genes with H3K9ac into five quantiles according to gene expression, a large subset of genes was observed with the least expressed genes showing the lowest levels of H3K9ac and the most highly expressed genes showing the highest levels of H3K9ac; these results suggest that H3K9ac positively correlates with gene expression (Figure S2). Surprisingly, when we observed subtle differences in the H3K9ac deposition, a small subset of genes without detectable expression showed H3K9ac distribution in the upstream region of the TSSs (Figure S2). Whether the distribution of H3K9ac in the upstream region of promoter is involved in gene silencing is unknown. The dataset further expectedly revealed that global H3K9ac density was increased along with leaf aging, similar to that in *Arabidopsis* (Brusslan et al. 2015).

**Figure 4.**
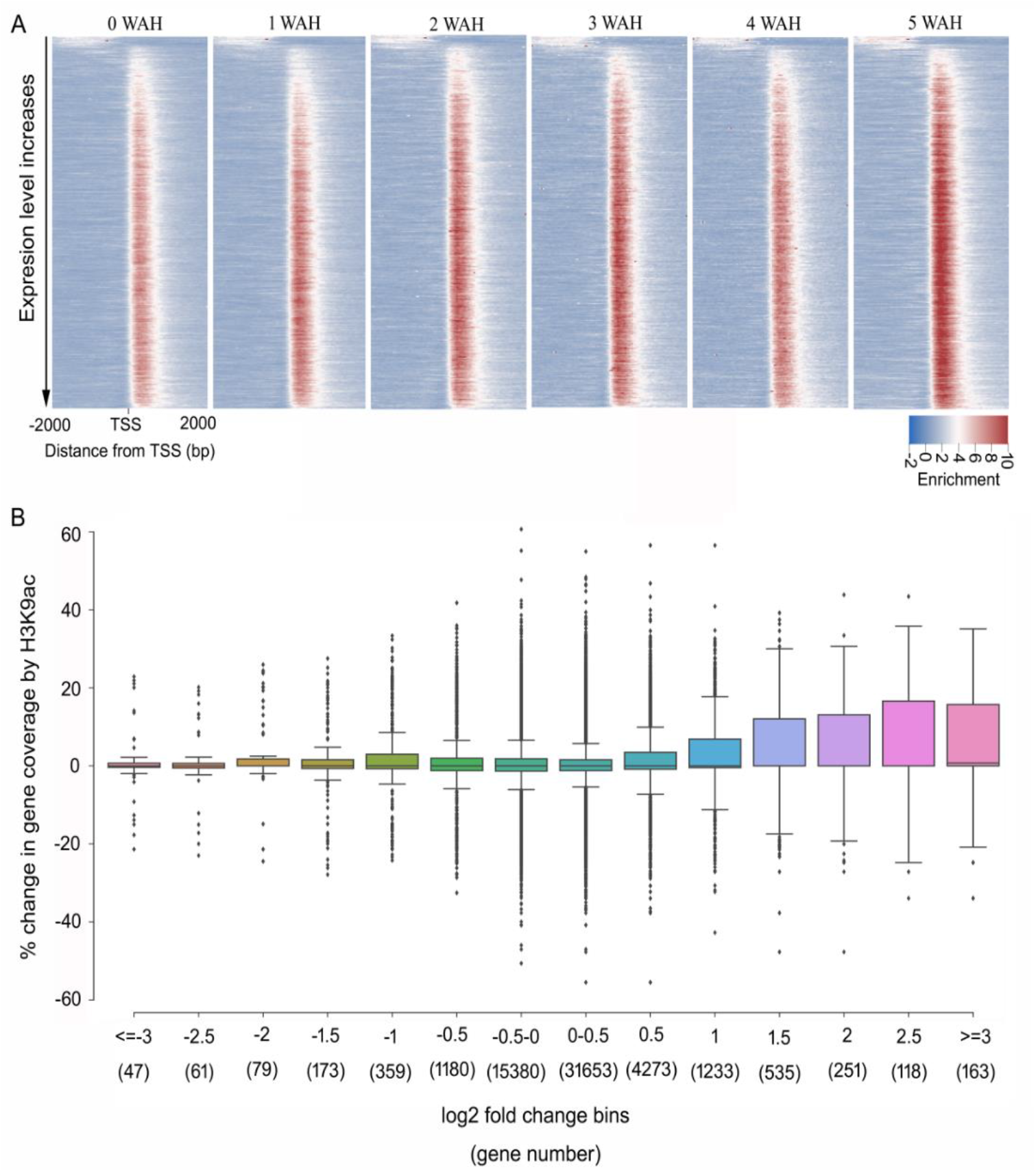
Relationship of the H3K9ac changes and gene expression during developmental flag senescence in rice. **(A)** Heat map of H3K9ac levels around the TSSs of all genes (from 2-kb upstream to 2-kb downstream regions of the TSSs). All genes marked with H3K9ac were sorted by their expression level measured by RNA-seq. Each row represents 50 genes, and each column represents the normalized H3K9ac enrichment level in each 40-bp bin. Red equates to increased signals and blue is the reverse. **(B)** Breadth of H3K9ac modification and gene expression during leaf senescence. The percentage change in H3K9ac gene coverage is plotted for gene expression bins. Genes are placed in bins according to the fold change in gene expression across five time points relative to 0 WAH. Gene numbers per bin are given in parentheses.

Except for the H3K9ac density, more interestingly, the peak width of the H3K9ac became broader with higher gene expression (Figure 4A). To better illustrate the association between the breadth of H3K9ac markers and the transcript levels of genes during leaf aging, genes were placed into differential bins based on their log2 fold change of transcript level at individual time points with respect to those at the initial time point (0 WAH). The change in the proportion of the H3K9ac marker coverage was determined across the corresponding comparison for each bin; their distributions are plotted in Figure 4B. The correlation analysis showed that up-regulated genes with the greatest fold change displayed the largest positively correlated change in coverage for H3K9ac. However, when the fold change in the down-regulated genes increased, the breadth of H3K9ac coverage remained at a similar level. These findings suggest that there is only a positive correlation between breadth of H3K9ac coverage and expression of up-regulated genes during rice flag leaf aging (Figure 4B, Figure S3).

To gain insight into the set of genes that showed increased expression as senescence correlated with more H3K9ac coverage, we performed a GO enrichment analysis. The biological process GO term ‘regulation of transcript, DNA-templated’ (GO: 0006355) was enriched in significantly up-regulated DEGs (Log2expression fold change>1) with broad H3K9ac peaks (Figure S4). The results suggest that the breadth of H3K9ac coverage may control transcriptional regulatory events. Interestingly, the breadth of H3K9ac coverage is extended to the gene body (Figure 4A). It is possible that the breadth of H3K9ac coverage may directly affect transcript elongation, which is supported by H3K9ac enrichment accompanying RNAPII recruitment and transcript elongation (Gates et al., 2017; Huang et al., 2018; Vaid et al., 2020). These results suggest that the density and peak width of H3K9ac plays important role in dictating the transcriptional activity and elongation of genes during rice flag leaf aging.

### Integrative analysis of ChIP-seq and RNA-seq DEGs identifies metabolic pathway associated genes with differential H3K9ac during rice flag leaf aging

H3K9ac is an active marker, and several SAGs, such as *AtWRKY53, AtERF6*, and *HvS40*, are activated in parallel with H3K9ac enrichment (Huang et al., 2018; Janack et al., 2016; Brusslan et al. 2015). To identify H3K9ac-associated DEG targets, we focused on a total of 2,719 genes that showed combined alteration in both gene expression and H3K9ac modification (Figure 5A; Supplementary Dataset S3). Among them, 46% (1,249 of 2,719) of up-regulated DEGs gained H3K9ac, whereas 37% (996 of 2,719) of down-regulated DEGs displayed a loss of H3K9ac marker (Figure 5A). The remaining 17% of H3K9ac-associated DEGs displayed down-regulated DEGs with increased H3K9ac and up-regulated DEGs with a loss of H3K9ac. This suggests a strong positive correlation between H3K9ac modification and gene expression during rice flag leaf aging.

**Figure 5.**
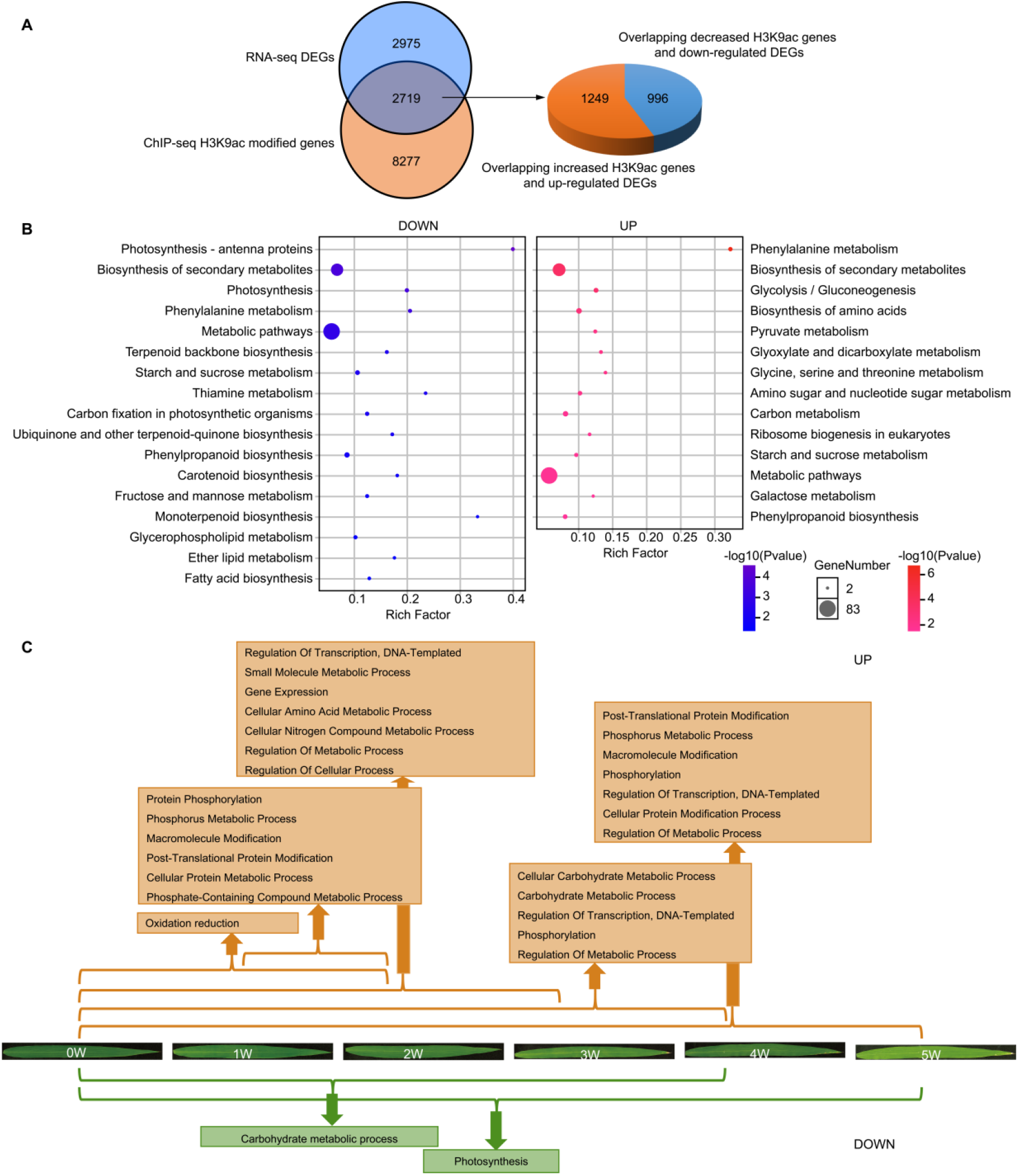
Integration of ChIP-seq and RNA-seq Data to identify metabolic -associated genes with differential H3K9ac levels during leaf aging. **(A)** Venn diagram showing the overlap of 5,694 DEGs from RNA-seq data and 10,996 differential H3K9ac–associated genes from ChIP-seq data. A set of 2,719 genes in the overlapping regions were identified including 1,249 up-regulated DEGs with increased H3K9ac levels and 996 down-regulated DEGs with decreased H3K9ac levels. **(B)** Functional analysis of genes significantly changing in both down-(left panel) or both up-regulated (right panel) H3K9ac modification and gene expression. The rich factor is the ratio of the number of DEGs annotated in a given pathway to the number of all genes annotated in the pathway. A higher rich factor indicates greater intensity. The P value ranges from 0 to 1, with a lower P value indicating greater intensity. The size of the circle indicates the number of genes. Blue indicates down regulated, Red indicates up regulated. **(C)** Key biological processes initiated or repressed of H3K9ac-associated differential display expression genes at different time points during rice flag leaf aging. Enriched GO terms were identified in groups of genes that significantly change in H3K9ac modification and gene expression during leaf development and senescence. Brown pale is both enriched and upregulated; green pale is both lose and down regulated.

KEGG pathway enrichment analysis revealed that 1,249 up-regulated H3K9ac-gained DEGs were mainly enriched in metabolic pathways, for instance, genes were significantly enriched in amino acid metabolism pathways (e.g. phenylalanine metabolism), and in carbohydrate metabolism pathways (e.g. glycolysis/ gluconeogenesis) (Figure 5B; Supplementary Dataset S4); 996 down-regulated H3K9ac-lose DEGs were mostly presented in energy metabolism pathways including photosynthesis and carbon fixation, others presented in small molecule metabolism pathways (e.g. biosynthesis of secondary metabolites, amino acid, carbohydrates and lipid metabolism pathways) (Figure 5B; Supplementary Dataset S4). These results suggest that H3K9ac modification may systemically influence the expression of genes associated with diverse metabolism during leaf development.

GO analysis of the H3K9ac-associated DEGs showed the key overrepresented biological processes. Enriched H3K9ac and up-regulated DEGs (hUP-gUP) genes were significantly related to oxidation reduction and protein phosphorylation in first two weeks, followed by regulation of transcription, small molecules (amino acid and nitrogen compound) metabolic processes, and regulation of metabolic processes at 3 WAH, followed by carbohydrate metabolic processes at 4 WAH, and finally cellular protein modification processes (phosphorylation). In contrast, lose H3K9ac and down-regulated DEGs (hDN-gDN) genes within 4 WAH were mostly related to the carbohydrate metabolic processes, followed by photosynthesis at 5 WAH (Figure 5C; Supplementary Dataset S5). Taken together, these data reflect an alteration in H3K9ac abundance as an important response in developmental and metabolism-related genes in rice flag leaf transition to senescence.

### Integrative analysis of ChIP-seq and RNA-seq DEGs identifies leaf senescence-associated genes with differential H3K9ac during leaf aging

A comparison of 2,719 H3K9ac-associated DEGs genes with 188 rice SAGs from LSD 3.0 (http://www.ricedata.cn/) database (Li et al., 2020), in addition to the current reported rice senescence-associated genes, has identified 36 rice DEG SAGs relative to H3K9ac (Figure 6A; Supplementary Dataset S3). These H3K9ac–associated senescence-related DEGs are shown in Figure 6B. We further compared 1,249 hUP-gUP and 996 hDN-gDN genes with 36 rice SAGs (Figure 6; Supplementary Dataset S3). Of them, 20 hUP-gUP SAGs included genes related to Chl degradation, such as *STAY GREEN RICE* gene (*OsSGR;* Jiang et al., 2007), *NON-YELLOW COLORING1* (Os*NYC1)*, Os*NYC3*, and senescence-activated genes, such as *RICE NAC PROTEIN* (*OsNAP), miR164-TARGETED NAC* gene (*OsOMTN4*; Fang et al., 2014), *cZ-O-GLUCOSYLTRANSFERASES* (*cZOGT1*; Kudo et al, 2012), *CONIFERALDEHYDE 5-HYDROXYLASE* (*OsCAld5H1*; Takeda et al., 2017), *GLYCOSYLTRANSFERASE (OsAkaGal), CYTOCHROME P450* gene (*CYP94C2b*; Kurotani et al., 2015), *PECTIN METHYL ESTERASE* (*OsPME1), DUF584 Family member 7(OsS40-7), OsMYC2*, and *OsWRKY14*, which were continuously upregulated, while 12 hDN-gDN SAGs included LOC_Os01g68450 encoding an unknown expressed protein, GLYCINE DECARBOXYLASE COMPLEX H-protein (*OsGDCH*; Lin et al., 2016), *PHYTOCHROME INTERACTING FACTOR-LIKE 1* (*OsPIFL1*; Todaka et al., 2012), *STAY GREEN RICE LIKE* (*OsSGRL), MADS-box protein 56 (OsMADS56)*, and *Lipoxygenase 2 (OsLOX2)*, which exhibited continual downregulation during leaf aging (Figure 6; Supplementary Dataset S3). Only four genes were oppositely hUP-gDN- or hDN-gUP-regulated, including GLYCOSYL TRANSFERASE (*OsPLS3*), THYLAKOID FORMATION1(*OsTF1*), HEAVY METAL-ASSOCIATED DOMAIN CONTAINING PROTEIN (*OsHMAD*), and CATALASE (*OsCATB*) (Figure 6; Supplementary Dataset S3). Our findings suggest that there is a significant regulatory role of H3K9ac with respect to the series of SAGs during leaf senescence.

**Figure 6.**
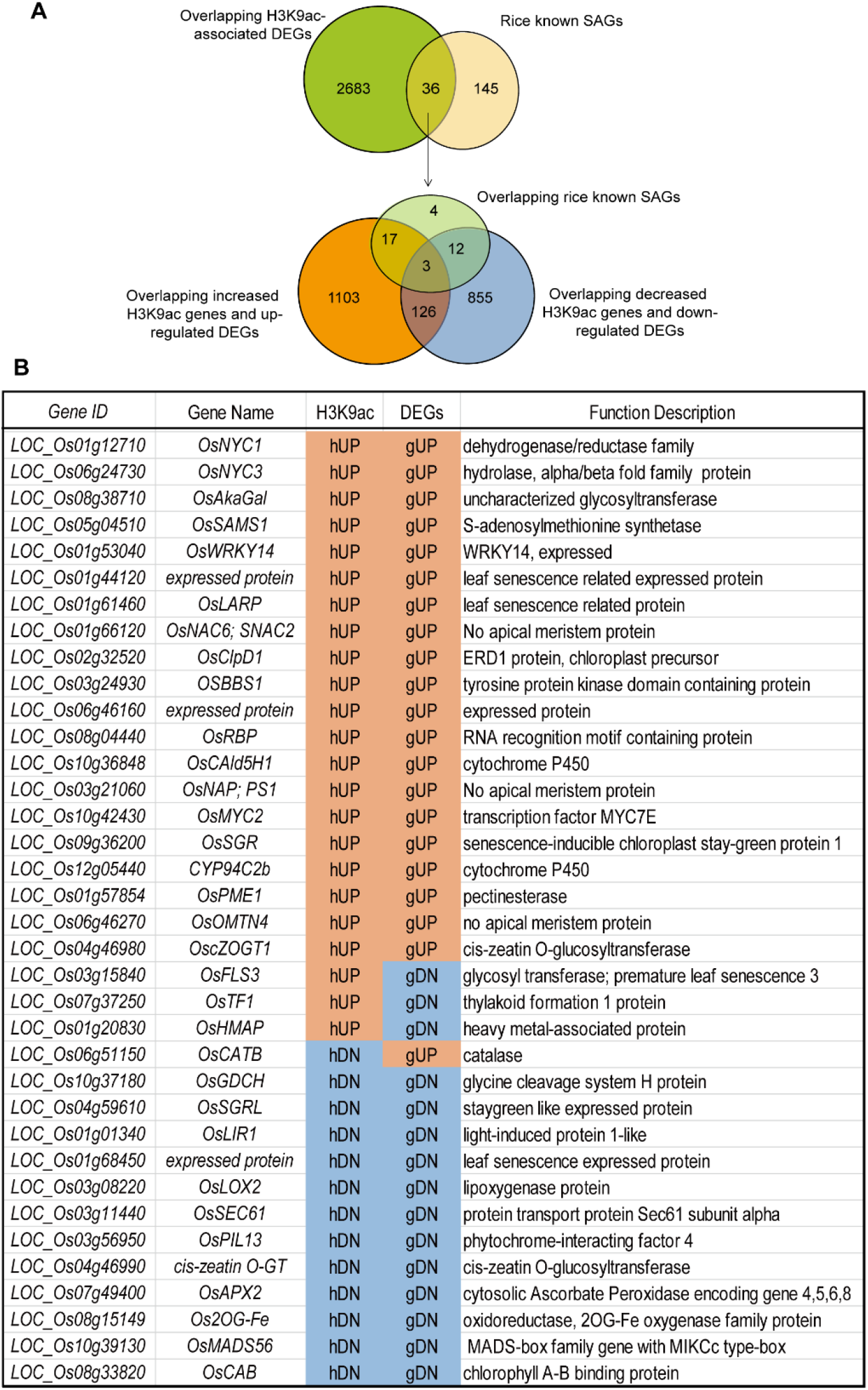
Integration of ChIP-seq and RNA-seq Data to identify leaf senescence-associated genes with differential H3K9ac levels. (A) Venn diagram showing the overlap SAGs from rice LSD 3.0 and 2,719 genes from Figure 5A. 36 H3K9ac-associated differential expression SAGs are identified during leaf aging. Among them, 20 genes are hUP-gUP, 12 genes are hDN-gDN, 4 genes are hUP-gDN or hDN-gUP. (B) The list of SAGs is illustrated their function description.

We further identified the H3K9ac-associated DEGs at each stage of leaf aging (Figure 7A). Strikingly, we found several SAGs such as *SGR, OMTN4, CYP94C2b, cZOGT1, OsCAld5H1*, and one expressed protein (*LOC_Os06g46160*) exhibit a significantly increased transcript level that accompanies enrichment of H3K9ac at a late stage of leaf aging (4 WAH and 5 WAH). In contrast, *OsGDCH* and *OsPIL1* displayed a continuous decline in both transcript and H3K9ac levels from an early stage (2 WAH) of leaf aging (Figure 7A; Supplementary Dataset 3), which negatively regulates leaf senescence in rice. hUP-gUP and hDN-gDN SAGs during leaf aging are shown in Figure 7B. H3K9ac and RNA-seq profiles of these known SAGs during leaf aging were visualized using Integrative Genomics Viewer (Figure S6 and Figure S7).

**Figure 7.**
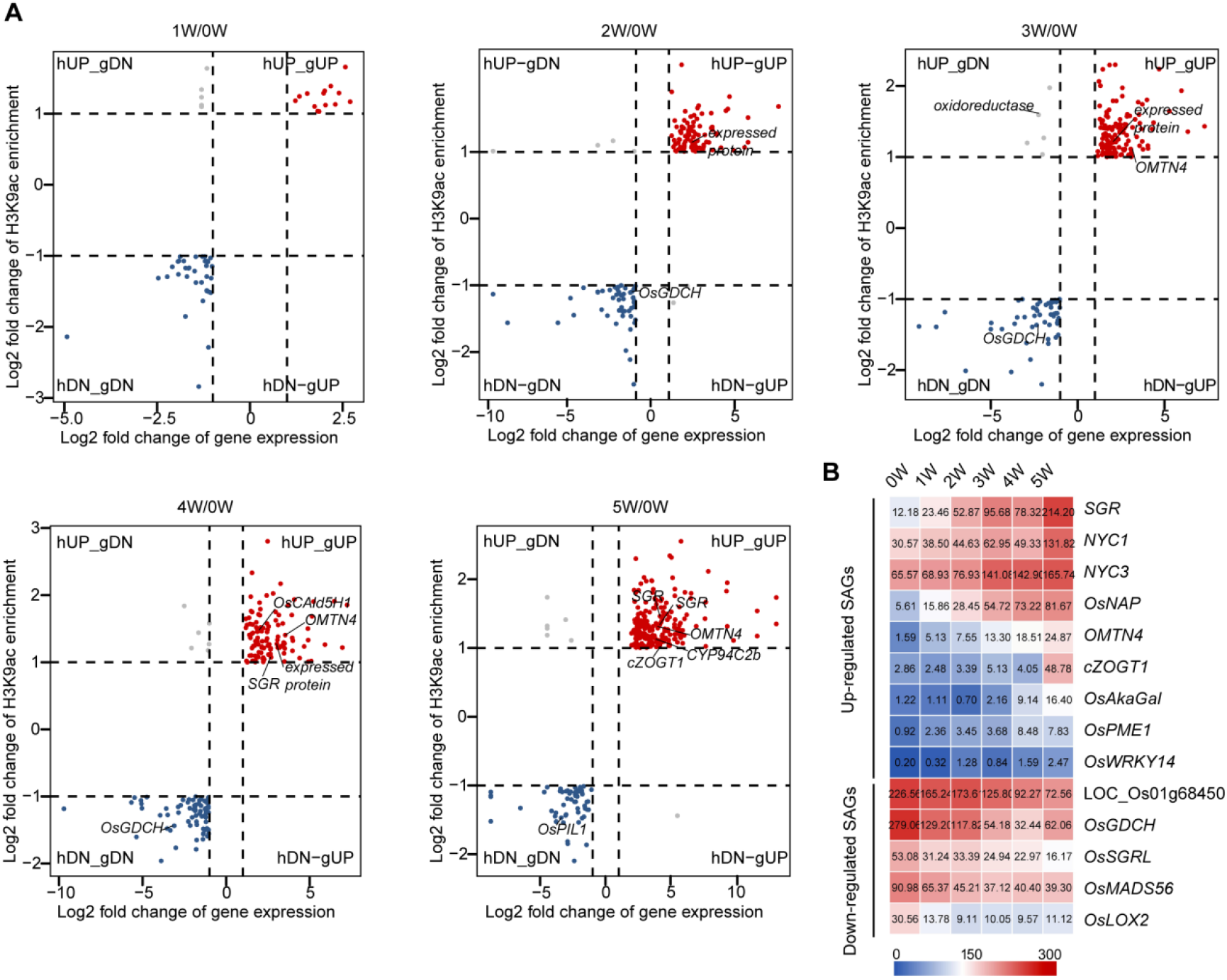
Alteration of gene expression and H3K9ac enrichment for each stage of leaf aging compared with 0W. Plots for log2 fold change of gene expression and H3K9ac enrichment for each stage of aging compared with 0W. Genes with a significant increase or decrease both in H3K9ac and gene expression are highlighted in red and blue, respectively. Note that genes may have more than one H3K9ac peak site, thus genes with a significant increase/decrease in H3K9ac. Inducing expression downregulation/ upregulation is highlighted in dark-gray. The horizontal and vertical dashed lines highlight a 2-fold increase or decrease. Rice known or putative SAGs with a significant increment or reduction of H3k9ac and gene expression are respectively labeled in each stage of aging versus 0W.

A representative single gene profile of H3K9ac modification and gene expression was further confirmed by chromatin immunoprecipitation, qPCR (ChIP-qPCR) and RT-qPCR. Genome tracks of RNA-seq and ChIP-seq data for hUP-gUP candidate genes, *S40-7* and *OsbHLH1*, are indicated in Figure 8A and Figure 8D. The transcript levels of *S40-7* and *OsbHLH1* were continuously increased during leaf aging (Figure 8B and Figure 8E); the enrichment levels of H3K9ac of *S40-7* and *OsbHLH1* were mostly increased from 2 WAH to 5 WAH, except for at 1 WAH (Figure 8C and F). In contrast, the hDN-gDN candidate gene *OsGDCH* (Figure 8G) and its transcript level and H3K9ac enrichment level clearly declined during leaf aging (Figure 8H and I). Expression patterns of nine differentially H3K9ac-modified genes during rice flag leaf senescence are shown in supplementary Figure (Figure S8). All confirmed a positive correlation between the change in H3K9ac enrichment and the change in senescence-related gene expression.

**Figure 8.**
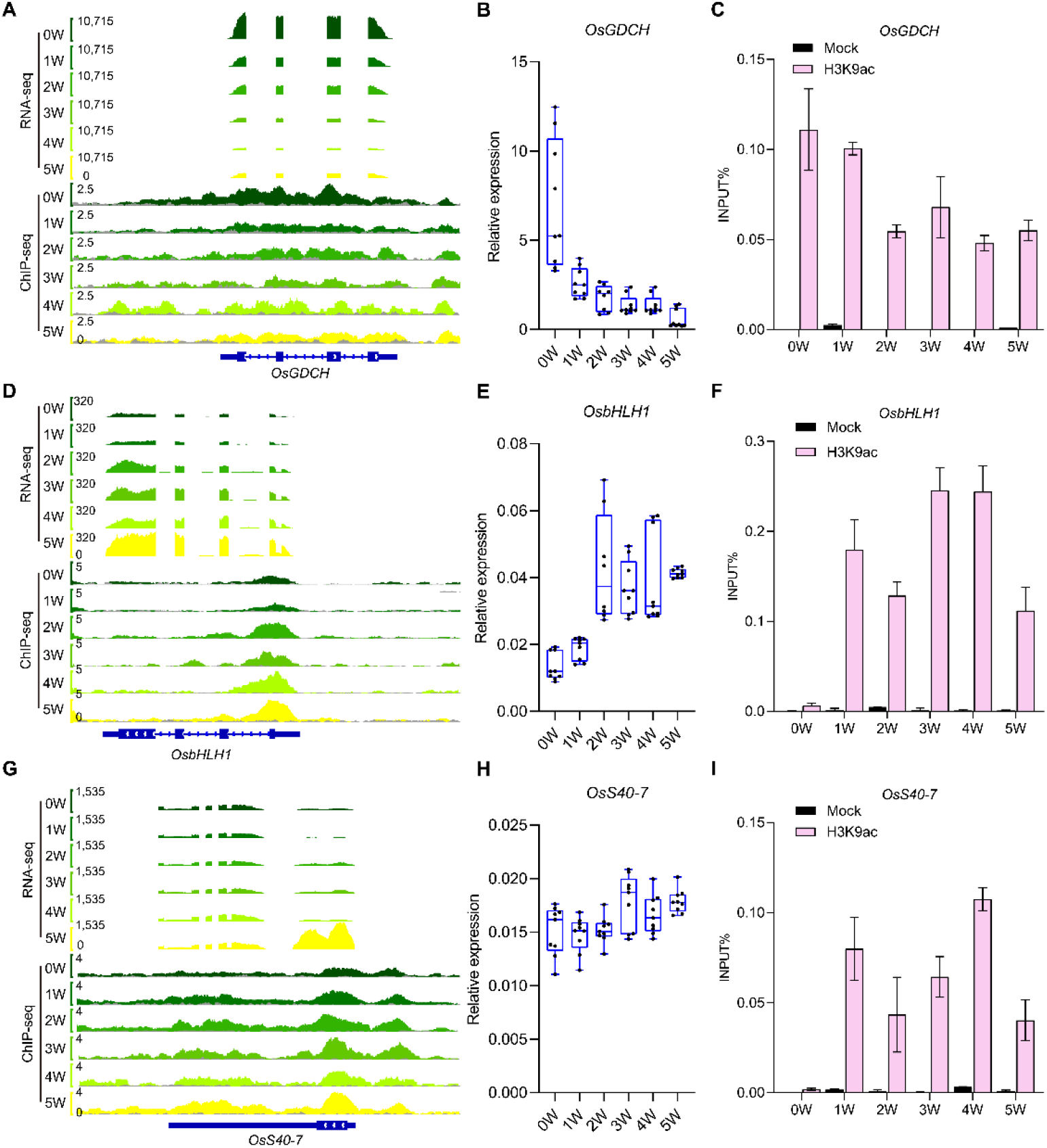
Representative single gene profiles of H3K9ac modification and gene expression were examined by ChIP-qPCR and RT-qPCR. **(A), (D), (G)** Genome tracks of RNA-seq and ChIP-seq data for *OsS40-7* (LOC_Os01g52730) (A), *OsbHLH1* (LOC_Os07g43530) (D), and *OsGDCH* (LOC_Os10g37180) (G) at different stages of leaf aging. Y-axis value means normalized read counts. Data from younger leaves are dark green, whereas data from older leaves are yellow. Input is in dark-gray. **(B), (E), (H)** RT-qPCR analysis for gene expression levels of *OsS40-7(B, OsbHLH1(E)*, and *OsGDCH(H)* at different stages of leaf senescence. *OsACTIN* was used as an internal reference. Error bars = ±SD (n = 3) of three biological replicates **(C), (F), (I)** ChIP-qPCR analysis of H3K9ac levels of *OsS40-7*(C), *OsbHLH1*(F) and *OsGDCH*(I) at different stages of leaf senescence. Error bars = ±SD (n = 3) of three biological replicates. ChIP enrichment was determined as % of input.

### Integrative analysis of ChIP-seq and RNA-seq DEGs identifies regulatory genes with differential H3K9ac

The MapMan analysis of the H3K9ac-associated DEGs showed that abundant genes were predominately involved in ‘signaling’ pathways, for instance, in hormone signaling pathways, including auxins, ABA, ethylene, SA, and JA, in protein kinases, cell wall, proteolysis, and secondary metabolite signaling; abundant genes were also TFs (Figure 9A; Supplementary Dataset S6).

**Figure 9.**
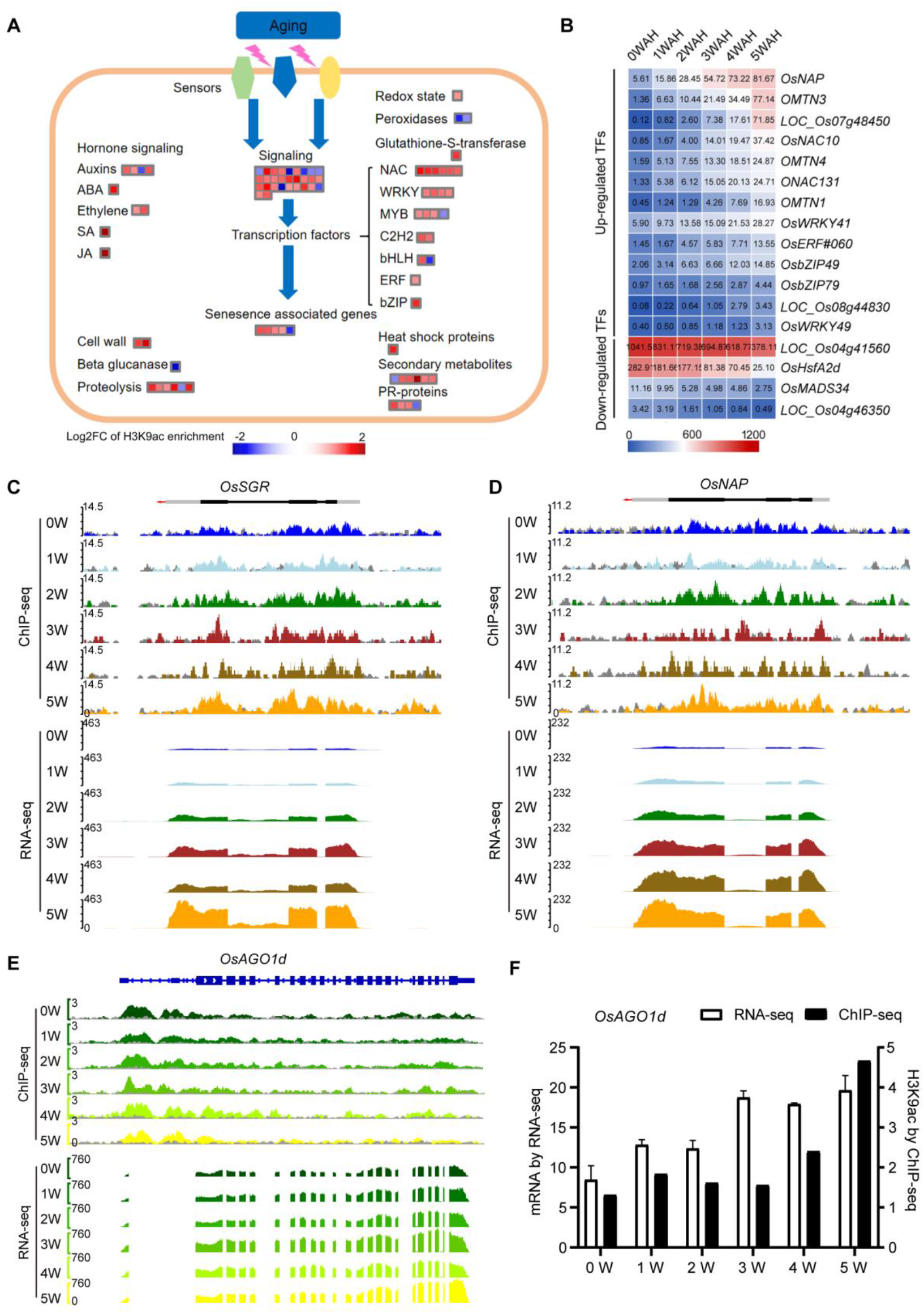
Integration of ChIP-seq and RNA-seq Data to identify regulatory genes with differential H3K9ac levels during leaf aging. (A) Assignment of differential H3K9 acetylated genes among 5WAH vs. 0 WAH during leaf aging in Mapman bins. The red and blue squares indicate the hyper- and hypoacetylated genes. (B) Expression profiles for the represented overlapping TFs during flag leaf aging. Red indicates a higher expression level, whereas blue indicates a lower expression level. (C) Genome tracks of ChIP-seq and RNA-seq data for *OsSGR* (LOC_Os09g36200) with a significant increase both in H3K9ac and gene expression at different leaf aging stages. *OsSGR* is an over-represented senescence associated gene derived from Figure 9A. Y axis value means normalized read counts. Input is in dark-gray. (D) Genome tracks of ChIP-seq and RNA-seq data for *OsNAP* (LOC_Os03g21060) with an increase both in H3K9ac and gene expression. *OsNAP* is an over-represented senescence associated TF gene derived from Figure 9B. Y axis value means normalized read counts. Input is in dark-gray. **(E)** Genome tracks of RNA-seq and ChIP-seq data for *OsAGO1d* (LOC_Os06g51310) at different leaf aging stages. Y-axis value means normalized read counts. Data from younger leaves are darker green, whereas data from older leaves are more yellow. Input is in dark-gray. **(F)** The gene expression level and H3K9ac level of *OsAGO1d* are shown in the barcharts. Left Y-axis denotes mRNA expression by RNA-seq. Right Y- axis denotes H3K9ac enrichment of *OsAGL1d* by ChIP-seq.

Hormone biosynthesis- and signaling-related genes were analyzed as previously described (Zhou et al., 2017). A total of 76 H3K9ac-associated phytohormones-related DEGs with 45 hUP-gUP and 38 hDN-gDN combinations have been observed during leaf aging. The result revealed that many H3K9ac-associated DEGs are key genes involved in ABA signaling and transport, ethylene biosynthesis and signaling, and jasmonic acid biosynthesis and signaling (Figure 9A; Supplementary Dataset S3). We also queried rice kinase genes in the Rice Kinase Phylogenomics Database (http://ricephylogenomics. ucdavis.edu/kinase/index. shtml). A total of 154 putative rice kinase genes were identified from the H3K9ac-associated DEGs. A set of 91 genes showed increased H3K9ac and upregulated gene expression, while 75 genes showed down-regulated expression accompanied by a loss in H3K9ac markers (Figure 9A; Figure 9C; Supplementary Dataset S3).

To characterize the H3K9ac-associated DEGs TFs, we queried rice TFs in the Plant Transcription Factor Database (http://planttfdb.cbi.pku.edu.cn/index.php). A total of 173 H3K9ac-associated TF DEGs with the 108 hUP-gUP and 80 hDN-gDN combinations have been identified, including NAC, WRKY, ERF, MYB and bHLH TF members (Supplementary Dataset S3). Most of the NAC, WRKY and ERF TFs (genes) were accompanied by increased H3K9ac markers and mRNA levels during leaf aging, including *OsNAP, OMTN4* and *OsWRKY14* mentioned above (Figure 7C; Figure 9B; Figure 9D).

We used the CYTOSCAPE program or KEGG to analyze all H3K9ac-associated DEGs and showed regulation of transcription was in the middle of a regulatory network, controlling diverse metabolism. Cellular biosynthesis and organic substance biosynthesis are the key notes of macromolecular biosynthesis (Figure 5B and Figure S4). Therefore, our results reveal a complex regulatory network of metabolism-mediated senescence associated with H3K9ac during leaf aging in rice.

### H3K9ac regulates non-coding RNA biogenesis during rice flag leaf aging

Several miRNAs, such as *miR164* and *miR319*, in *Arabidopsis*, maize and rice have been reported to be involved in regulating leaf senescence (Kim et al., 2009; Schommer et al., 2008; Thatcher et al., 2015; Qin et al., 2016). A previous study suggested that histone acetylation may also regulate miRNA expression at different levels (Kim et al., 2009; Pulido and Laufs, et al., 2010; Lee at al., 2011). We speculated that H3K9ac likely affects these components in the non-coding RNA biosynthesis pathway. To test this possibility, eight Dicer-like (*OsDCLs*), 19 Argonatute (*OsAGOs*) and five RNA-dependent RNA polymerase (*OsRDRs*) genes in rice (Kapoor et al., 2008) were collected to linked analysis with the H3K9ac actyltome and transcriptome. Notably, results showed that among 32 genes, 27 genes gained the H3K9ac marker, while five genes were associated with loss of the H3K9ac marker; there was no change in gene expression level observed during leaf aging with the exception of *OsAGO1d*, which coordinately altered H3K9ac enrichment and gene expression (Figure 9E-F). This result suggests that H3K9ac may be associated with non-coding RNAs, but is likely directly related to microRNA biosynthesis as *AGO1* determines microRNA biosynthesis in plants (Vaucheret, 2008; Suarze et al., 2015; Wu et al., 2020).

## DISCUSSION

It has been shown that H3K9ac dynamics is closely associated with gene expression alteration during plant development and adaptation to the changing environment (Pan et al., 2017; Zheng et al., 2019; Li et al., 2019). H3K9ac has been implicated in leaf senescence in *Arabidopsis* (Brusslan et al., 2015; Hinckley et al., 2019). Although transcription of a few genes of rice were previously reported to be associated with to H3K9ac (Huang et al, 2007; Zhong et al., 2013; Fang et al., 2016), it is worth noting a global landscape of H3K9 acetylation-modified gene expression targets during flag leaf aging. Rice flag leaf aging, a slow, extent leaf aging period including nutrition redistribution and allocation, directly related to grain filling, which is a fine-tunes aging period, it also is much better material for studying on leaf senescence (flag leaf, 8 weeks) (Schippers et al., 2015). In this study, a global analysis of histone acetylatome and transcriptome revealed that H3K9ac enrichment activates gene transcription and transcript elongation. We established a landscape of H3K9 acetylation-modified gene expression targets during flag leaf aging, including many known metabolism-related enzyme genes, senescence-associated TFs and signal transduction-related genes, as well as miRNA biosynthesis-related genes, such as *AGO1* during flag leaf aging. Our findings reveal a complex regulatory network and metabolism-mediated senescence network associated with H3K9ac in rice.

### Temporal dynamics of H3K9ac at specific loci related to gene transcription and elongation during flag leaf aging in rice

The conserved distribution pattern of H3K9ac at the TSSs both in rice (this study) and in *Arabidopsis* (Brusslan et al., 2015) suggests that H3K9ac plays a significant role in gene transcription regulation during leaf aging. This study further shows a coordinated regulation of H3K9ac enrichment and gene activation, in which genes with low levels of H3K9ac enrichment showed low levels of expression, while genes with high levels of H3K9ac enrichment exhibited high levels of expression, when H3K9ac was positioned after the TSSs (Figure 4A; Figure S2). However, a subset of genes with no expression showed H3K9ac distribution in the upstream region of the TSSs (Figure S2). This could be due to involvement of H3K9ac in alterative histone modification, such as methylation (Huang et al., 2018), ubiquitination (Lan et al., 2018) or miRNA co-repression (Kim et al., 2009; Pulido and Laufs, et al., 2010; Lee et al., 2011) in the upstream promoter, which could affect downstream gene silencing. In order to better understand this, the potential involvement of H3K9ac in gene silencing will need to be further investigated.

Our study additionally revealed that global H3K9ac density was increasing with leaf aging, and the peak width of the H3K9ac modification was broader around genes with higher expression (Figure 4A). It is consistent with the breadth of histone markers correlating with gene expression during *Arabidopsis* aging (Brusslan et al., 2015; You et al., 2017). This also suggests that there is a positive correlation between density and breadth of H3K9ac with expression of up-regulated genes during rice flag leaf aging (Figure 4B, Figure S3). Regulation of transcription was enriched for the up-regulated genes with the broadest H3K9ac coverage, suggesting that H3K9ac coverage may control regulatory loci. Interestingly, the breadth of H3K9ac coverage is extended to the gene body (Figure 4A). In *Mammalian* and *Arabidopsis*, as well as rice, the breadth of H3K9ac coverage is also extended to the gene body (Brusslan et al., 2015; Du et al., 2013; Gates et al., 2017). Our results suggest that the breadth of H3K9ac coverage may affect transcript elongation, in a conserved manner. It has reported that H3K9ac enrichment accompanies RNAPII recruitment and transcript elongation at the promoter region of the downstream target gene in *Arabidopsis* cells, mammalian *Hela* cells, mouse *ESCs* cells, and *Drosophila* S2 cells (Gates et al., 2017; Huang et al., 2018; Vaid et al., 2020; Etchegaray et al., 2020). In *Hela* cells, H3K9ac specifically recruits specific reader proteins for transcription progression and recruitment of the super elongation complex (SEC) may help direct progression of transcription into elongation. H3K9ac helps to designate the transition from transcription initiation to elongation on chromatin (Vermeulen et al., 2010; Bian et al., 2011). Therefore, these results support the notion that in rice, the density and peak width of H3K9ac may play an important role in dictating the transcriptional activity and elongation of genes during flag leaf aging. The detailed mechanism can be addressed by using release of promoter–proximal paused Pol II in response to histone acetylation of its downstream gene body in the future.

### The reprogramming of gene expression during flag leaf aging is under epigenetic control

Reprogramming of epigenetic states is critical for the initial establishment and subsequent maintenance of lineage-specific transcriptional programs (You et al., 2017; Li et al., 2020). Using the leaf senescence model, material of monocot (rice) flag leaf aging, our integrated transcriptomic and epigenomic analyses show a landscape of H3K9ac-associated DEGs in a fine-tuned manner. The hDN-gDN genes within 4 WAH are mostly related to the carbohydrate metabolic process due to a grain filling stage, followed by photosynthesis at 5WAH, showing a yellowing or chlorophyll degradation phenotype (Figure 1). In contrast, the hUP-gUP genes were significantly enriched for genes related to oxidation reduction, followed by regulation of transcription, amino acid small molecules metabolic processes, and finally cellular protein modification processes (phosphorylation) (Figure 5C) from 0 WAH to 5 WAH, showing active nutrient scavenging and remobilization (Lim et al., 2007). The small set of hUP-gDN or hDN-gUP genes, such as gene related to THYLAKOID FORMATION 1, which is involved in thylakoid formation and variegated leaves; this is not a typical senescence protein (Wang et al., 2004; Wangdi et al., 2010). In addition, PLS3, HMAD, and OsCATB, which are all have a major role in pathogen response, were involved in detoxification or defense during leaf aging (Yu et al., 2016; Pradhan et al., 2017; Wang et al., 2018). Taken together, this profiling highlights alteration in H3K9ac abundance as being an important response in the regulation of metabolism-related genes in rice flag leaf transition to senescence.

The 2,149 hUP-gUP and 996 hDN-gDN sets of genes are most likely to represent the direct effects of differential H3K9ac enrichments on gene expression. For example, (1) SAGs: *SGR, OMTN4, CYP94C2b*, and *cZOGT1* (Figure 6B), which are related to metabolism of photosynthesis carbohydrate (Jiang et al., 2007; Fang et al., 2014; Kurotani et al., 2015; Kudo et al., 2012), exhibit a significantly increased expression level accompanied by increasing H3K9ac during rice flag leaf aging. (2) TFs: 173 identified TFs, including 26 WRKY TF genes, 19 NAC genes, 20 ERF and 22 bHLH genes (Supplementary Dataset S2). Many of these genes are known to play important roles in regulating leaf senescence. Examples include *OsNAP, OsNAC6, OsPIL16, OsMYC2, OsWRKY14, OsMADS56, OsPIL1*, and *OsS40-7* (Figure 7C), which control nitrogen, ion, photosynthesis carbohydrate, and secondary metabolite biosynthesis (Liang et al., 2014; Lee et al., 2017; Yang et al., 2018; Lim et al., 2020; Li et al., 2021; Sakuraba et al., 2017; Habiba et al., 2021). (3) Signal transduction and phytohormones-related genes. For instance, a receptor kinase of rice OsBBS1/OsRLCK109, the *bbs1* mutant, exhibited hypersensitivity to salt stress and early leaf senescence phenotypes (Zeng et al., 2018). In addition, 76 H3K9ac-associated phytohormones-related DEGs have been observed during flag leaf aging. This result reveals that many H3K9ac-modified DEGs are key TFs and metabolism-related enzymes with ABA signaling and transport, ethylene biosynthesis and signaling, and especially jasmonic acid biosynthesis and signaling and their crosstalk (Supplemental Dataset S3). Genes such as *OsPME1, OsTSD2, OsMYC2, CYP94C2b*, and *OsLOX2* are all involved in JA biosynthesis and signaling (Fang et al. 2016); *cZOGT1* and *cZOGT2* are involved in rice cytokinin metabolism, and overexpression of *cZOGT1* and *cZOGT2* genes has been shown to lead to short-shoot phenotypes, delay of leaf senescence, and a decrease in crown root number (Kudo et al., 2012). (4) Interestingly, OsAGO1d coordinately alters H3K9ac enrichment and gene expression during flag leaf aging (Figure 9). It has already been established that AGO1 determines microRNA biosynthesis and plant development in *Arabidopsis* and rice (Vaucheret, 2008; Suarze et al., 2015; Li et al., 2019; Wu et al., 2020). Thus, it is possible that H3K9ac may be associated with microRNA biosynthesis. Recent studies have shown that in tomato, H3K4me3 and H3K9ac markers, and RNAPII enriched in the SlyWRKY75 intronic region, where this miRNA has been predicted to bind, and the presence of histone markers and RNAPII on the structural gene of Sly-miR1127-3p miRNA in plants occur in response to pathogen infection (Crespo-Salvador et al., 2020). In *Maize* and *Arabidopsis*, it has been previously described that the endogenous siRNA biogenesis pathway requires RNA-dependent RNA polymerase-2 (RDR2). A loss-of-function of RDR2 caused dramatic reduction of 24-nucleotide siRNAs, however, slight increase of the three activating epigenetic markers, H3K4me3, H3K9ac, and H3K36me3, in roots relative to shoots of *Arabidopsis* and *Maize*, respectively (Wang et al., 2009). Moreover, examples of posttranscriptional regulation mediated by small RNAs have been identified during leaf senescence (Kim et al., 2009; Thatcher et al., 2015) and may also explain some of the inconsistencies between H3K9ac marks and gene expression. We will speculate on the correlation between H3K9 acetylome and miRNAome. Thus, numerous important primary regulators of senescence do display a concomitant change in H3K9ac markers and gene expression.

It is not surprising that previous studies have shown that many histone acetyltransferases and histone deacetylases are involved in leaf senescence in *Arabidopsis*. For example, AtHDA19 is a negative regulator of leaf senescence (Tian and Chen, 2001), while AtHDA6 and AtHDA9 are positive regulators of leaf senescence (Wu et al., 2008; Chen et al., 2016). In rice, there are 9 histone acetyltransferases and 18 histone deacetylases (Banerjee and Roychoudhury, 2017). OsSRT1, a rice homolog of SIR2 (SILENT INFORMATION REGULATOR2), could deacetylate H3K9ac and alter the expression of genes related to programmed cell death and aging, and negatively regulate leaf senescence. (Zhong et al, 2013). Recently, it has reported that over-expressing OsHDA710 delays leaf senescence and participates in ABA signaling (Zhao et al. 2020; Ullah et al. 2021). The further insight into the function of histone acetyltransferases and histone deacetylases will be helpful to illustrate the mechanism of epigenetically controlling leaf senescence.

Besides, a pool of three replicates of IPs and inputs of each developmental stage flag leaves was used to run ChIP-seq. Although the analysis of dataset from one pool of three biological replicates for ChIP-seq existed somewhat limitation of the study, fifteen senescence associated genes randomly selected were confirmed by using an independent batch of ChIP DNA for ChIP-qPCR and RT-qPCR assay, the results that are similar to those seen by ChIP-seq and RNA-seq (Figure 8) illustrated a good repeatability of our ChIP-seq data in a certain extent.

## Acknowledgements

This work was supported by the grant of National Natural Science Foundation of China (grant number 32001437 to Y.Z., 31801266 to Y.K.X., 31770318 to Y.M.), and by the grant of National Science Foundation of Fujian Province (grant number2021J02025 to Y.M.).

## Author contributions

Y.M. designed the study. Y.Z. performed the data analysis. Y.Y.Z., Y.Y.L. and Y.Z. carried out ChIP-seq, RNA-seq, ChIP-qPCR and RT-qPCR experiments. Z.Y. and Z.Y.Z. performed the integrative analysis and L.F.G. supervised the analysis. D.Y.Z. and X.N.W. performed the WGCNA analysis. B.F.L. and D.D.H. performed RT-qPCR. Y.Z., Y.K.X. and Y.Y.L. provided technical assistance to all. Y.Z. and Y.M. wrote the article. All authors edited the article.

## Conflicts of Interest

The authors declare no conflicts of interest.

## Supplementary data

**Figure S1**. Characteristics of rice flag leaves during aging.

**Figure S2**. Relationship of normalized, average H3K9ac density to different gene expression levels.

**Figure S3**. Relationship of the gene coverage of H3K9ac to gene expression levels.

**Figure S4**. Cytoscape exhibited a biological network of H3K9ac-enriched DEGs.

**Figure S5**. Plots for Log2 fold change of gene expression and H3K9ac enrichment for different stage comparisons.

**Figure S6**. Genome tracks of ChIP-seq and RNA-seq data for six over-represented rice known genes with a significant increase both in H3K9ac and gene expression at different leaf aging stages.

**Figure S7**. Genome tracks of ChIP-seq and RNA-seq data for two SAGs (*OsNYC1*, LOC_Os01g12710; *OsNYC3*, LOC_Os06g24730) with an increase and two over-represented rice known genes (*OsPIL1*, LOC_Os03g56950; *oxidoreductase*, LOC_Os08g15149) with a significant decrease both in H3K9ac and gene expression. Y axis value means normalized read counts. Input is in dark-gray.

**Figure S8**. Expression patterns of differential H3K9ac-modified genes during flag leaf senescence in rice.

**Table S1**. Read and alignment summary for ChIP-seq libraries. **Table S2**. Distribution of H3K9ac peaks in different gene structures. **Table S3**. The list of primer pairs used in this study.

**Supplementary Dataset S1**. The total enriched KEGG pathways for differential H3K9ac-modified genes during rice developmental flag leaf aging.

**Supplementary Dataset S2**. DEGs in each stage-specific module.

**Supplementary Dataset S3**. Integrative analysis overlapping genes of ChIP-seq and RNA-seq DEGs, identifying TF genes, protein kinase, hormone-related genes, rice known SAGs.

**Supplementary Dataset S4**. KEGG enrichment analysis of 1,249 both up- and 996 both-down regulated H3Kac and expression level genes.

**Supplementary Dataset S5**. GO enrichment analysis of 1,249 both up- and 996 both-down regulated H3Kac and expression level genes.

**Supplementary Dataset S6**. Integrative Analysis of ChIP-seq and RNA-seq Data for 5 WAH vs 0 WAH.

## Notes

### Competing Interest Statement

The authors have declared no competing interest.

